# Location of CD39+ T cell sub-populations within tumours predict differential outcomes in non-small cell lung cancer

**DOI:** 10.1101/2022.09.29.509921

**Authors:** Lilian Koppensteiner, Layla Mathieson, Samuel Pattle, David A Dorward, Richard O’Connor, Ahsan Akram

## Abstract

An improved mechanistic understanding of immunosuppressive pathways in NSCLC is important to develop novel diagnostic and therapeutic approaches. Here, we reveal that the prognostic significance of the rate limiting ectonucleotidases in adenosine production CD39 and CD73 requires knowledge of cell type specific expression and localisation within tumours. In a cohort of early treatment naïve NSCLC patients, high stromal expression of CD39 and CD73 predicts poor outcome. CD39 expression amongst T cells identifies CD39+CD4+ Tregs which predict poor outcome and CD39+CD103+ CTL which confer a survival benefit if high densities are observed inside of the tumour nest. Bulk RNA Seq shows that the TME of NSCLC upregulates regulatory pathways in CD4+ T cells and exhaustion in CD8+ T cells. Analysis of single-cell RNASeq datasets illustrates that CD39+CD4+ Tregs are enriched in Treg signature gene sets, and CD39+CD103+ CTL show gene signatures indicative of an exhausted cytotoxic phenotype with an upregulated expression of CXCL13.

Combined knowledge of patterns of distribution and location are required to understand the prognostic impact of CD39+ T cell populations in NSCLC. This study provides an improved understanding of the spatial and functional characteristics of CD39+ T cells and illustrates their significance to patient outcome.

## Introduction

The immune response to cancer depends on efficient cytotoxic T lymphocytes (CTL) and checkpoint inhibition therapy is aimed at reinvigorating local cytotoxic T cells. However, the rate of immunotherapy failure in non-small cell lung cancer (NSCLC) is as high as 80% in unselected patients.^1^ Immunogenic hot tumours with high densities of tumour infiltrating lymphocytes are linked to improved response to immunotherapy.^2^ Therefore identifying the extent of cytotoxic lymphocyte infiltration, as a measure of a pre-existing immunity could be used to predict clinical outcome or responsiveness to treatment. In recent years, CD39 has emerged as a marker to identify tumour relevant T cells. Unlike PD1, CD39 expression is lacking in bystander CD8+ T cells and highly expressed in CD8+ T cells specific for tumour antigens in colorectal cancer and melanoma.^3,4^ CD39 expression on CTL also describes a terminally exhausted population in viral infections and tumours. ^3–7^

Duhen et al showed that CD39^+^CD8^+^TIL often co-express the tissue resident memory (TRM) marker CD103.^8^ In head and neck squamous cell cancer (HNSCC) and ovarian cancer, these CD39^+^CD103^+^CD8^+^ cells display an exhausted tissue resident memory T cell phenotype that, despite its reduced capacity for effector cytokine production, shows great cytotoxicity against neoplastic cells and can be found across human solid malignancies.^8^ Functionally, CD39 is an outer membrane enzyme that converts extracellular ATP (eATP) into adenosine monophosphate (AMP), which is then hydrolysed into adenosine by CD73. Adenosine has highly immunosuppressive effects, as it enhances the activity of suppressive immune cells including tumour associated macrophages, myeloid derives suppressor cells and regulatory T cells (Tregs), and inhibits neutrophils, NK cells, DCs and T cells, which ultimately results in tumour growth and progression.^9^ In order to investigate the distribution of adenosine generating ectonucleosides we characterized CD39, CD73 and CD103 in NSCLC via flow cytometry and multiplex Immunofluorescence and investigated their functional relevance using a publicly available NSCLC single cell RNA sequencing dataset.^10^ Patients with high CD39 and CD73 expression in close proximity to each other show markedly worse outcome. CD39+CD4+ T cells which upregulate genes associated with an active Treg phenotype (CD25) also predicts poor outcome while high densities of CD39+CD103+CD8+ T cells in the tumour nest which upregulate cytotoxicity - and tissue residency genes, are linked to improved 5-year survival.

## Methods

### Ethics statement

Healthy volunteer blood was obtained following informed consent and the study was approved by Lothian Regional Ethics Committee (REC) (REC No: 20-HV-069) prior to enrolment in the studies. Tumour/non-cancerous lung tissue and blood samples were obtained following approval by NHS Lothian REC and facilitated by NHS Lothian SAHSC Bioresource (REC No: 15/ES/0094), with informed consent from all patients. The tissue microarray was approved NHS Lothian REC and facilitated by NHS Lothian SAHSC Bioresource (REC No: 15/ES/0094) and approved by delegated authority granted to R&D by the NHS Lothian Caldicott Guardian (Application number CRD19031)

### NSCLC tissue digest

NSCLC were identified by the pathologist with paried tumour and non-cancerous lung tissue taken from the same lobe of lung in fresh curative cancer resections. As previously described in O’Connor et. al, samples were mechanically minced, spun at 350g for 5 minutes, supernatant removed and samples digested with 1mg/ml Collagenase IV (Merck) and 1mg/ml DNAse (Merck) in Media for 1 hour at 37°C (with vortexing every 10 minutes). Digested samples were passed through 70μm filters, followed by centrifugation at 350g for 5 minutes. Supernatants were removed and red blood cells were lysed with RBC lysis buffer (Biolegend). Cells were washed again, counted, and stained for Flow cytometry.

### Peripheral blood mononuclear cell (PBMC) extraction

Blood samples from early-stage non-small cell lung cancer patients were paired to respective tissue samples. Healthy donor blood samples were donated by healthy volunteers at the Centre for inflammation Research in the Queen’s Medical Research Institute in Edinburgh. PBMCs were isolated from whole blood using lymphoprep (Stemcell technologies) and SepMate tubes for density gradient centrifugation (Stemcell technologies). Cells were washed, centrifuged at 350g for 5 minutes, and stained for Flow cytometry.

### Flow cytometry

Cells were washed with 2ml of serum free DPBS and spun at 350g for 5 minutes. The supernatant was removed, and this step was repeated. Cells were stained with Zombie Live Dead UV (Biolegend) (1μl per test in PBS) to label dead cells and incubated for 30 minutes at room temperature in the dark. The reaction was stopped by adding 2 mls FACS buffer (DPBS +2% FCS) to each tube, followed by centrifugation at 350g for 5 minutes and removal of the supernatant. Prior to surface marker staining, cells were treated with FC block for 5-10 minutes at room temperature, to reduce non-specific binding (5μl/ test in 50μl FACS buffer). Antibodies were prepared at the required concentration in FACS buffer to a final volume of 50μl/test. Antibody panels are illustrated in Table 1. Antibody cocktails were added, and cells were incubated for 20 minutes at 4°C. 2ml FACS buffer was added to each tube to stop the reaction, followed by centrifugation at 350g for 5 minutes, after which cells were resuspended in 150μl fixation buffer (Biolegend) and 150μl FACS buffer. Samples were stored at 4°C in the dark until flow cytometry collection.

FACS data analysis was performed using FlowJo software. Gating strategy of tissue digest analysis is shown in supplementary Figure S1.

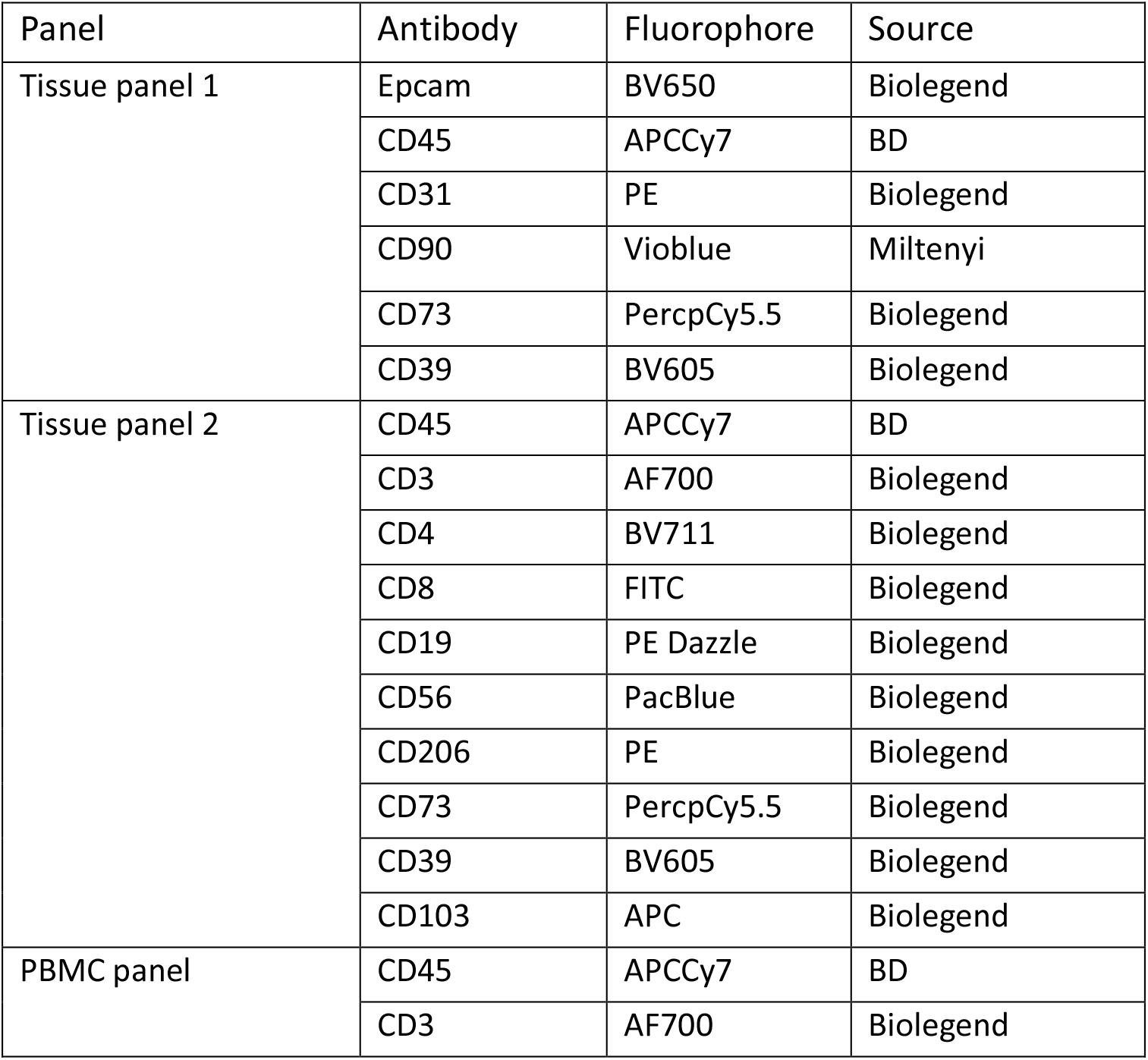
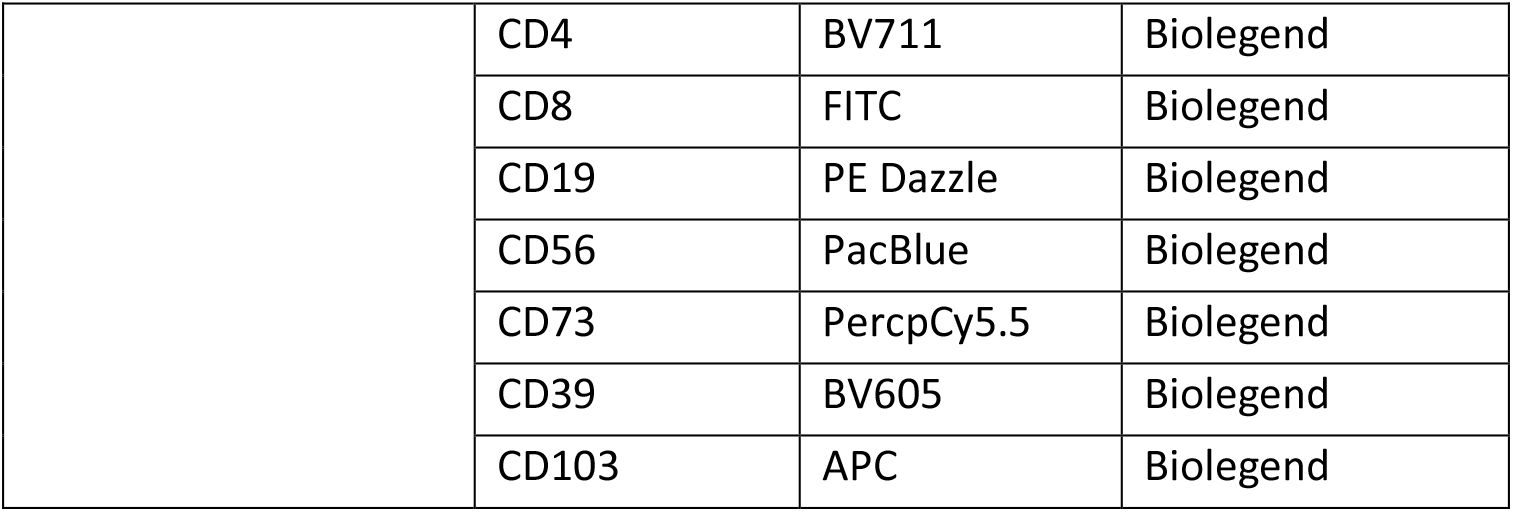

### Multiplex Immunofluorescence staining

A tissue microarray (TMA) was constructed from consecutive patients undergoing curative resection surgery for NSCLC in a regional thoracic centre. An experienced pathologist annotated each resection block and 1mm cores were taken and embedded into the TMA. All patients were followed up to assess for relapse and death. 4 μm sections were cut and slides were deparaffinised in Xylene and rehydrated in a series of ethanol dilutions. The following steps were performed on the Leica Bond automated staining robot. After initial heat-induced antigen retrieval (HIER) of 30min at 100°C, tissue slides were exposed to multiple staining cycles each including a 30 minute incubation with a protein block (Akoya), 1 hour incubation with the respective primary antibody, 30 minute incubation with the secondary antibody (Akoya), 10 minute incubation with the respective OPAL (Akoya) followed by 20 minute incubation with AR6 buffer (Akoya) at 85°C prior to the next staining cycles and finally stained with fluorescent DAPI (Akoya) for 10 minutes. In between each step, slides were washed with bond wash for 5 minutes.

### Multiplex Immunofluorescence imaging

The appropriate exposure time for image acquisition was set for each fluorophore by autoexposing on multiple (5-10) tissue areas per batch. Following fluorescence whole slide scans, regions of interest were selected for multispectral imaging (MSI) at 20x magnification, as scanning at 40x magnification is time-consuming and does not improve image analysis.^11^

### Multiplex Immunofluorescence analysis

MSI images were unmixed using representative snapshots of spectral library slides imaged at the same magnification and autofluorescence was isolated in InForm software. Unmixed images were exported and analysed in Qupath^12^ (Fig. S2A). Cell detection was performed using StarDist based on a watershed deep-learning algorithm and fluorescent threshold of DAPI nuclear staining.^13^ Following this, phenotyping was performed in a non-hierarchical manner by creating a composite classifier of single channel classifiers for each stain based on a fluorescent threshold that was adjusted for images if necessary. Ultimately, a machine learning algorithm was trained on multiple images to detect tumour and stroma areas. (Fig.S1B). Exported measurements included area measurements of tissue region and counts of cells that are positive for any combination of markers within these. Quantitative analysis was performed in Rstudio.

### Nanostring nCounter

NCL and tumour tissue samples from 6 patients were minced and digested as described above. Cells were stained for flow cytometry as described above using the following antibody panel:

| Antibody | Fluorophore | Source |
| --- | --- | --- |
| CD45 | APCCy7 | BD |
| CD3 | AF700 | Biolegend |
| CD4 | PE | Biolegend |
| CD8 | APC | Biolegend |
| CD11b | FITC | Biolegend |

CD4+ and CD8+ T cells were collected on the FACS Aria Fusion into RNAse free eppendorfs (Qiagen), transported on ice and cell pellets were resuspended and lysed in RLT buffer (Qiagen) prior to freezing at −80C. Nanostring nCounter of the CAR T cell panel was performed by Alison Munro at HTPU Institute of Genetics and Cancer Edinburgh. Only samples that passed nSolver QC were used for downstream analysis. Data from two separate nCounter runs was normalised in nSolver (Nanostring) using a calibration sample and normalized to HK genes selected using GeNorm algorithm. Normalised gene counts were used for Advanced Analysis in nSolver. Heatmaps of z-score of normalized counts of DEG (log2FC>/<0.5, p.value<0.05) were created using the pheatmap() function in R.

### Single cell RNA Sequencing data analysis

Single cell RNA sequencing data of T cells from human NSCLC tissue samples was downloaded from SCope (aertslab.org).^10^ CD4+, CD8+, CD39+ and CD103+ T cells were defined by gene expression counts of CD4 >0, CD8a >0, ENTPD1 >0 and ITGAE >0, respectively. Differential expression analysis was performed in R using the DESeq2 package.^14^ Volcano plots were created using the EnhancedVolcano() function. GO biological processes analyses of upregulated genes in CD39+ CD4+ T cells (log2Fold change >1, padj. <0.1) and CD39+CD103+CD8+ T cells (log2Fold change >0.5, padj. <0.1) were performed using the enrichGO() function of the R package clusterProfiler (OrgDb = org.Hs.eg.db, ont = BP). Gene set enrichment analysis (GSEA) of DEGs was performed using the gseGO() function of the R package clusterProfiler using gene sets obtained from Gene ontology biological processes database.

### TCGA data analysis

TCGA-LUAD and TCGA-LUSC datasets were used to analyse CD39 and CD73 expression and correlation to outcome. Pre-processed gene expression data (fragments per kilobase per million fragments, upper quartile normalized) of primary solid tumours and corresponding normal solid tissue as well as corresponding patient clinical data were downloaded using the GDC data transfer tool. High/ low expression of each marker was determined by the surv_cutpoint() function of the R survival package. Survival analysis was adjusted for sex, age, stage and smoking (pack years) via Cox multivariate regression analysis using the coxph() function in R.

## Results

### The TME of NSCLC exhibits upregulated expression of CD39

We first analysed the expression of CD39, CD73 and CD103 in peripheral blood and non-cancerous lung and tumour tissue of early untreated NSCLC using flow cytometry. (Fig.1A) Peripheral lymphocyte CD39 expression was limited to CD19+ cells, while low levels of CD39 expression were observed amongst CD4+ T cells, CD8+ T cells and CD56+ cells. We find an increase of CD39+ cells in non-cancerous lung (NCL) tissue compared with levels in peripheral blood, which was further elevated in the tumour microenvironment of NSCLC amongst T cells, in line with published reports of multiple human solid tumours.^7,15–19^ We additionally observed an upregulation of CD39 expression on CD56+ cells and CD90+ cells in tumour compared to NCL tissue. CD206+ cells are equally CD39+ in NCL and tumour tissue. (Fig.1B).

**Figure 1.**
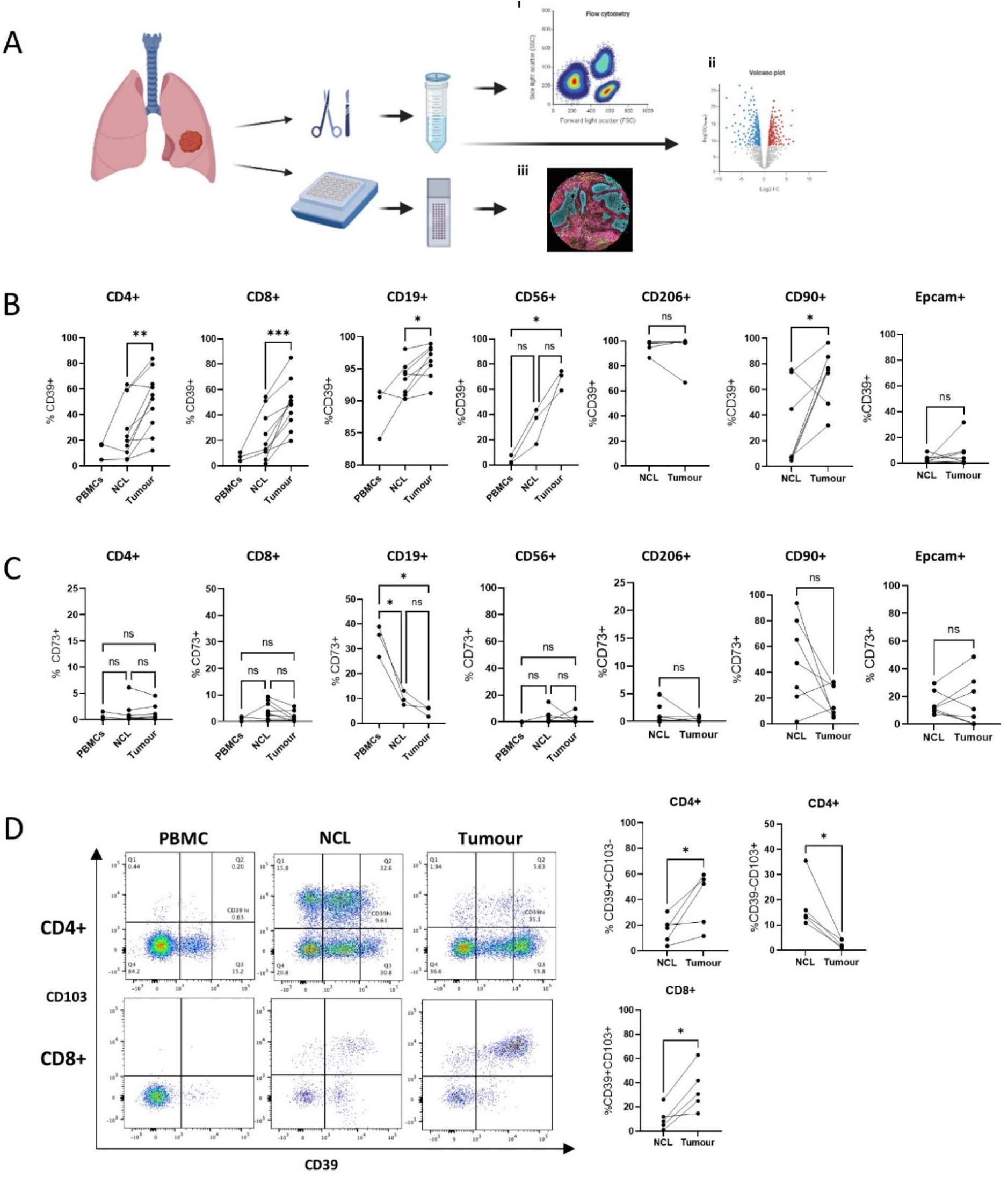
The TME of NSCLC exhibits upregulated expression of CD39. (A) Study design. (B) Expression levels of CD39 shown as % positive within subpopulations of CD4+, CD8+, CD19+ and CD56+ cells from peripheral blood, NCL and tumour tissue and CD206+, CD90+ and Epcam+ cells from NCL and tumour tissue of early stage NSCLC patients. (C) Expression levels of CD73 shown as % positive within subpopulations of CD4+, CD8+, CD19+ and CD56+ cells from peripheral blood, NCL and tumour tissue and CD206+, CD90+ and Epcam+ cells from NCL and tumour tissue of early NSCLC patients. (D) Representative staining of CD4+ and CD8+ T cells from a paired PBMC, NCL and tumour sample. Frequency of CD39+CD103- and CD39-CD103+ CD4+ T cells and CD39+CD103+ CD8+ T cells. Paired t tests were used for statistical analysis of 2 groups, one way ANOVAs with Tukeys multiple comparisons were used for comparing 3 groups. (* p=<0.05, ** p=<0.01, ***p=<0.001). Images created with Biorender.com.

Considering the downstream processing of CD39 synthesized cAMP into adenosine by CD73, we also analysed CD73 expression in NSCLC to further investigate which cell types contribute to adenosine synthesis in the TME of NSCLC. We observed high frequencies of CD73 expression amongst CD19+ cells and CD8+ T cells in the peripheral blood of healthy people which was significantly reduced in the peripheral blood of early NSCLC patients (Fig. S3) and further reduced in NCL and tumour tissue (Fig. 1C). Notably, some patients exhibited high levels of CD73 expression in CD90+ of NCL tissue, which was reduced in cancerous tissue. EPCAM+ cells display similar levels of CD73 expression in both NCL and tumour tissue (Fig.1C). While CD39 and CD73 expression is often associated with Tregs,^20^ our data illustrates that a variety of cell types harbour the potential to contribute to the immunosuppressive pressure of adenosine production via CD39 and these are enriched in the cancerous tissue. In stark contrast, we did not observe an upregulation of CD73 in the TME within cell types tested and saw highest frequencies amongst fibroblasts and EPCAM+ cells, indicating that adenosine production is likely potentiated by the presence of multiple cell types.

As previously shown ^8,21^ the majority of CD39+ CD8+ tumour infiltrating lymphocytes (TILS) co-express the tissue resident memory marker CD103. Conversely, amongst CD4+ T cells, we do observe a CD103+CD39-, as well as a CD103+CD39+ population in non-cancerous lung tissue, however, these are depleted in the cancerous tissue, where we see a shift towards a single positive CD39high population (Fig.1D). CD19+ cells, CD56+ cells and CD206+ cells display low frequencies of CD103, suggesting CD103 is exclusive to T cells in the lung tissue (Fig. S4).

### CD39, CD103 and CD73 display distinct spatial signatures in NSCLC

To assess the localisation of surface markers involved in adenosine production as well as T cell subpopulations characterised by CD39 and CD103 expression, we investigated their presence and spatial organisation in FFPE tissue of 162 early treatment naive NSCLC patients. CD39 expression was locally elevated in the tumour tissue compared to NCL (Fig.2A, B) where the vast majority was expressed in the tumour stroma (Fig.2C) on stromal cells, vascular structures and immune infiltrates (Fig.2D). Only some of these are CD4+ or CD8+ T cells confirming the presence of other CD39+ immune cells as shown by flow cytometry (Fig.2E). Conversely, CD103 was expressed in both tumour and stroma areas (Fig.2C) and mainly expressed by CD8+ T cells (Fig.2E), some of which displayed a CD39+CD103+ phenotype.

**Figure 2.**
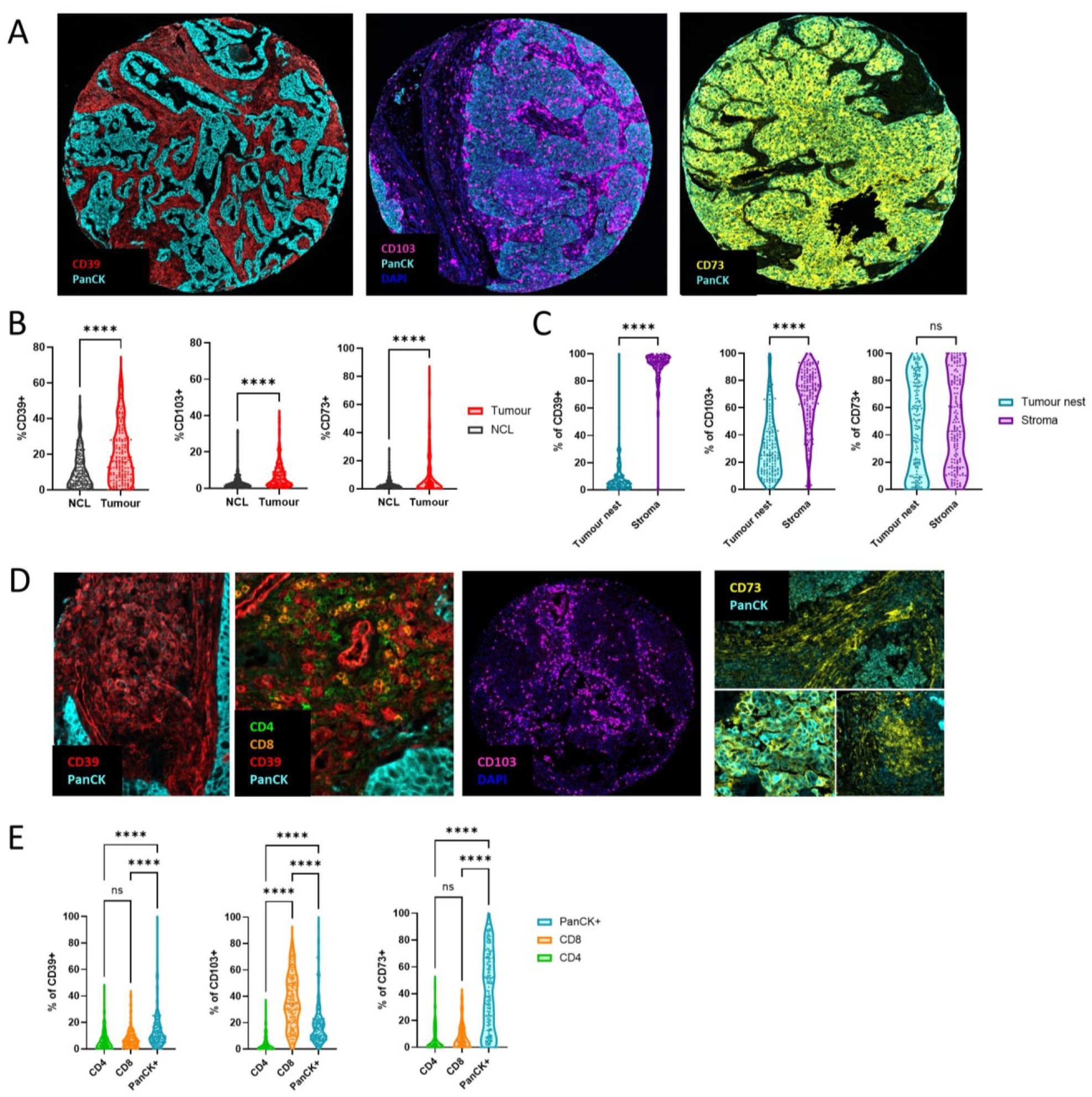
CD39, CD103 and CD73 display distinct spatial signatures in NSCLC. (A) Representative staining patterns of CD39, CD103 and CD73 in tumour tissue samples. (B) Frequency of CD39, CD103 and CD73+ cells in NCL and tumour tissue samples. (C) Distribution of CD39+, CD103+ and CD73+ cells across tumour nest and stroma. (D) Representative cell staining patterns of CD39, CD103 and CD73 in tumour tissue samples. (E) Distribution of CD39+, CD103+ and CD73+ cells across celltypes PanCK+, CD4+ and CD8+ cells. Unpaired t tests were used to compare NCL and Tumour groups. Paired t tests were used to compare tumour nest and stroma groups. Two-way ANOVAs were used to compare distribution across celltypes. (* p=<0.05, ** p=<0.01, ***p=<0.001)

CD73 expression was found in both tumour area staining up to 100% of cancer cells (Fig.2A) and in the tumour stroma (Fig.2C) staining T cell, non-T cell immune cells and fibroblasts (Fig.2D), demonstrating that co-localisation of CD39 and CD73 likely occurs in the tumour stroma.

### CD103-CD8+ T cells are unable to infiltrate the tumour nest

At cancerous tissue sites, the level of T cell infiltration was markedly elevated compared to NCL tissue (Fig. S5A), however, the majority of CD4+ and CD8+ T cells were confined to stromal areas (Fig.4A). While a higher frequency of CD39SP CD8+ T cells was found in the stroma, the frequency of CD39+CD103+ CD8+ T cells was equal in tumour nest and stroma and notably, the highest frequencies of CD103SP CD8+ T cells were observed inside of the tumour nest. (Fig. 4B,C) Immunofluorescence staining patterns illustrates that CD8+ T cells in the tumour nest are typically CD103+ and indeed, CD8+ T cells that did not express CD103 were restricted to the stromal area (Fig.4D, E) suggesting that CD103+ T cells are in closest proximity to cancer cells and CD103-CD8+ T cells either do not have the ability to infiltrate or that surrounding cancer cells induce CD103 upregulation in the tumour nest upon infiltration.

**Figure 3.**
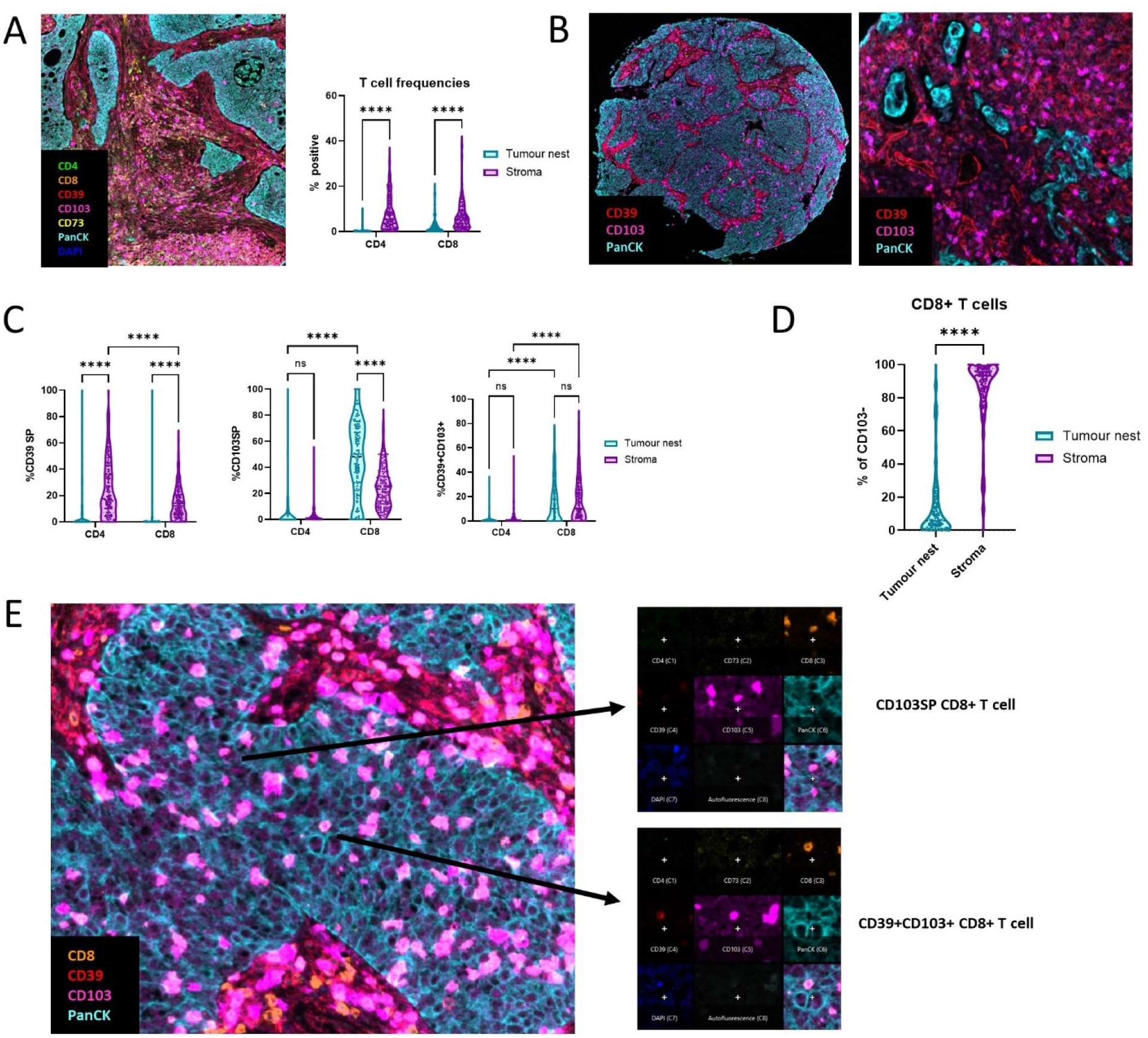
Infiltration of T cell subsets into the tumour nest of NSCLC. (A) Representative image of T cell restriction to the stroma and frequency of CD4+ and CD8+ T cells in tumour nest and stroma. (B) Representative images showing CD103 and CD39 expression. (C) Frequency of CD39SP, CD103SP and CD39+CD103+ cells within CD4+ and CD8+ T cells in tumour nest and stroma. (D) Distribution of CD103 negative CD8+ T cells across tumour nest and stroma. (E) Example image showing a CD103SP CD8+ TIL (top) and a CD39+CD103+ CD8+ TIL (bottom) in all 8 isolated fluorescent channels. Two-way ANOVAs were used to compare groups. (* p=<0.05, ** p=<0.01, ***p=<0.001)

**Figure 4.**
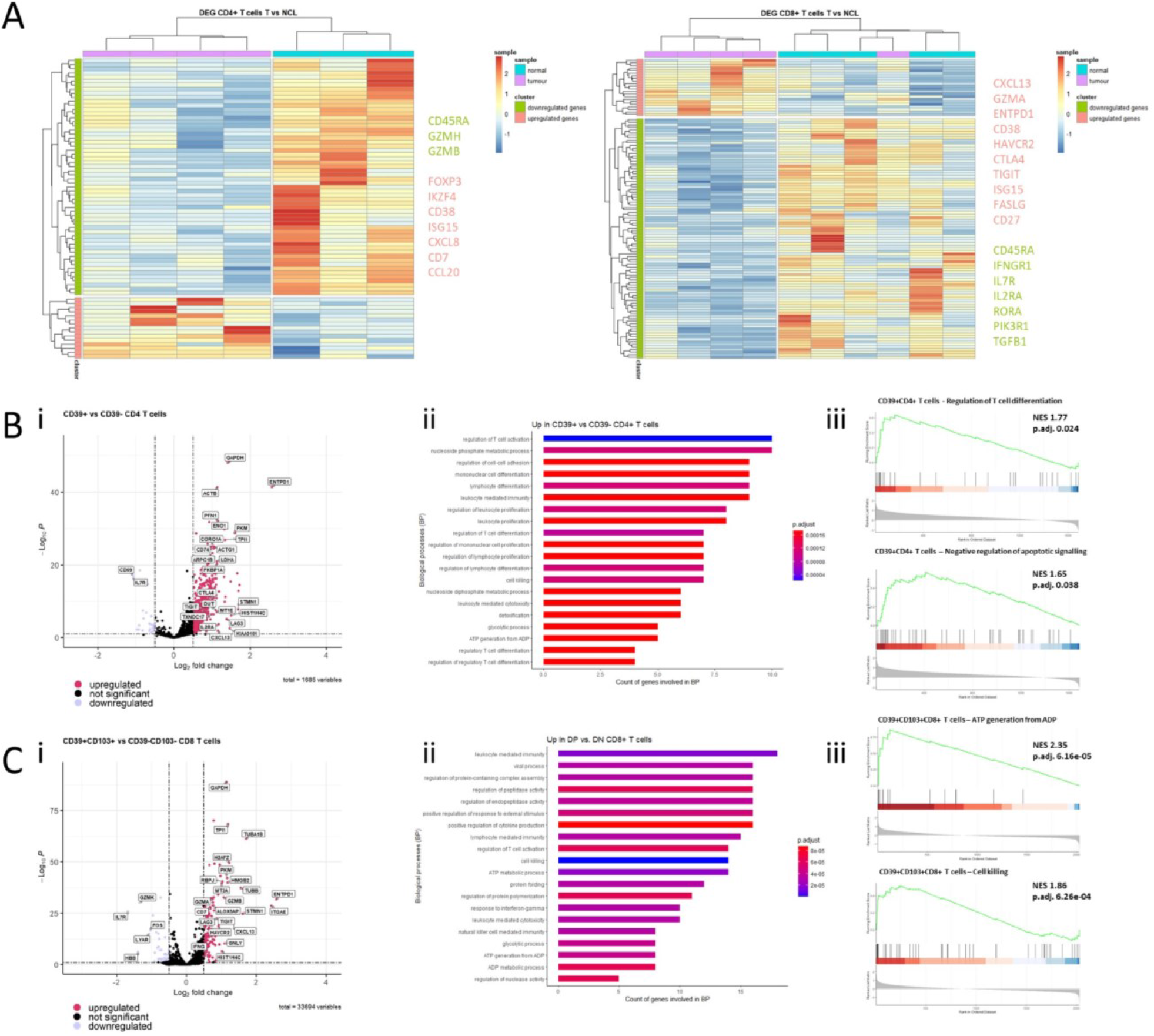
Transcriptomic and functional analysis of T cell subsets in NSCLC. (A) Heat maps showing unsupervised clustering of DEGs from Nanostring gene expression data of CD4+ (left) and CD8+ (right) T cells isolated from digested NCL and tumour tissue. (B) (i) Volcano plot showing DEG between CD39+ and CD39-CD4+ T cells. (ii) Bar plots of enrichment p values and gene count of respective gene sets of top 20 GO biological processes DEG in CD39+ CD4+ T cells compared to CD39-CD4+ T cells. (iii) GSEA of DEG in CD39+ CD4+ T cells compared to CD39-CD4+ T cells in regulation of T cell differentiation gene set (GO:0045580) and negative regulation of apoptotic signalling gene set (GO:2001234) (C) (i) Volcano plot showing DEG between CD39-CD103- and CD39+CD103+ CD8+ T cells. (ii) Bar plots of enrichment p values and gene count of respective gene sets of top 20 GO biological processes upregulated in CD39-CD103+ CD8+ T cells compared to CD39-CD103-CD8+ T cells. (iii) GSEA of DEG in CD39+CD103+CD8+ T cells compared to CD39-CD103-CD8+ T cells in ATP generation from ADP (GO:0006757) and cell killing (GO:0001906) gene sets.

### CD39+ T cells upregulate pathways of T cell exhaustion

We next performed Nanostring nCounter gene expression analysis of sorted CD4+ and CD8+ T cells from NCL and tumour tissue from early untreated NSCLC patients to analyse the effect of the TME on the T cell transcriptome. CD4+ T cells isolated from tumour tissue downregulate genes associated with activation (e.g. CD45RA) as well as granzymes B and H and upregulate genes associated with regulatory T cells (FOXP3, IKZF4, CD38, ISG15). Notably, CD8+ T cells downregulate activation and metabolism genes (CD45RA, IFNGR1, IL7R, IL2RA, RORA, PIK3R1) indicating the suppressive effect of the TME on CD8+ T cell function. Furthermore, the CD39 encoding gene ENTPD1 is amongst the most highly upregulated genes in CD8+ T cells isolated from tumour tissue, along with other T cell exhaustion marker encoding genes (HAVCR2, CTLA4, TIGIT). (Fig. 4A)

To further investigate the functional role of CD39+CD4+ and CD39+CD103+ CD8+ T cell subsets, we used a publicly available dataset ^10^ of single-cell RNA sequencing of NSCLC tumour tissue. CD39+CD4+ T cells showed downregulation of CD69 and IL7R expression, in line with reduced activation, and upregulation of genes associated with exhaustion (CTLA4, LAG3, TIGIT) (Fig.4Bi). Both Gene ontology (GO) and gene set enrichment analysis (GSEA) show a significant enrichment in gene sets of regulation of T cell activation-, differentiation, - proliferation and negative regulation of apoptotic signalling pathway genes further illustrating that CD39+CD4+ T cells represent a particularly suppressive regulatory T cell population (Fig.4Bii-iii).

In comparison to CD39-CD103-CD8+ T cells, CD39+CD103+ CD8+ T cells downregulate IL7R and the early activation marker CD69, and upregulate exhaustion markers (LAG3, TIGIT, HAVCR2), however, we also see an upregulated expression of IFNG and granzyme A and B indicative of effector function and high cytotoxicity. (Fig.4Ci) GO and GSEA analysis indicate a link to leucocyte mediated cytotoxicity, cell killing, and genes associated with ATP generation. (Fig.4Cii-iii)

Interestingly, both CD39+ populations (Fig.4B, C) show upregulated expression of the tissue resident marker ALOX5AP as well as CXCL13 expression, which has been described to be a feature of exhausted TILS and CD103+ TILS in cancer^15,22^, co-occurring with B cell recruitment and beneficial tertiary lymphoid structure (TLS) formation, in line with our flow data showing increased B cell frequencies in the TME of NSCLC. (Fig. S6)

### Spatial patterns of distinct T cell populations predict outcome in NSCLC

We hypothesized that CD39 and CD73, as enzymes responsible for adenosine synthesis, could negatively affect patient outcome. While overall expression of CD39 in our mxIF dataset does not affect recurrence-free survival (RFS), frequency of CD39+ stromal cells predicts reduced RFS probability at 5 years. Overall expression of CD73 has a negative predictive effect which is more pronounced when CD73 is expressed in the stroma. (Fig.5Ai) Moreover, a combination of high stromal CD39 and CD73 predicts reduced RFS probability suggesting that close proximity of CD39 and CD73 in the stroma could elevate locally produced adenosine levels, mediating strong immunosuppressive signalling around immune cells trapped in this area. (Fig.5B)

**Figure 5.**
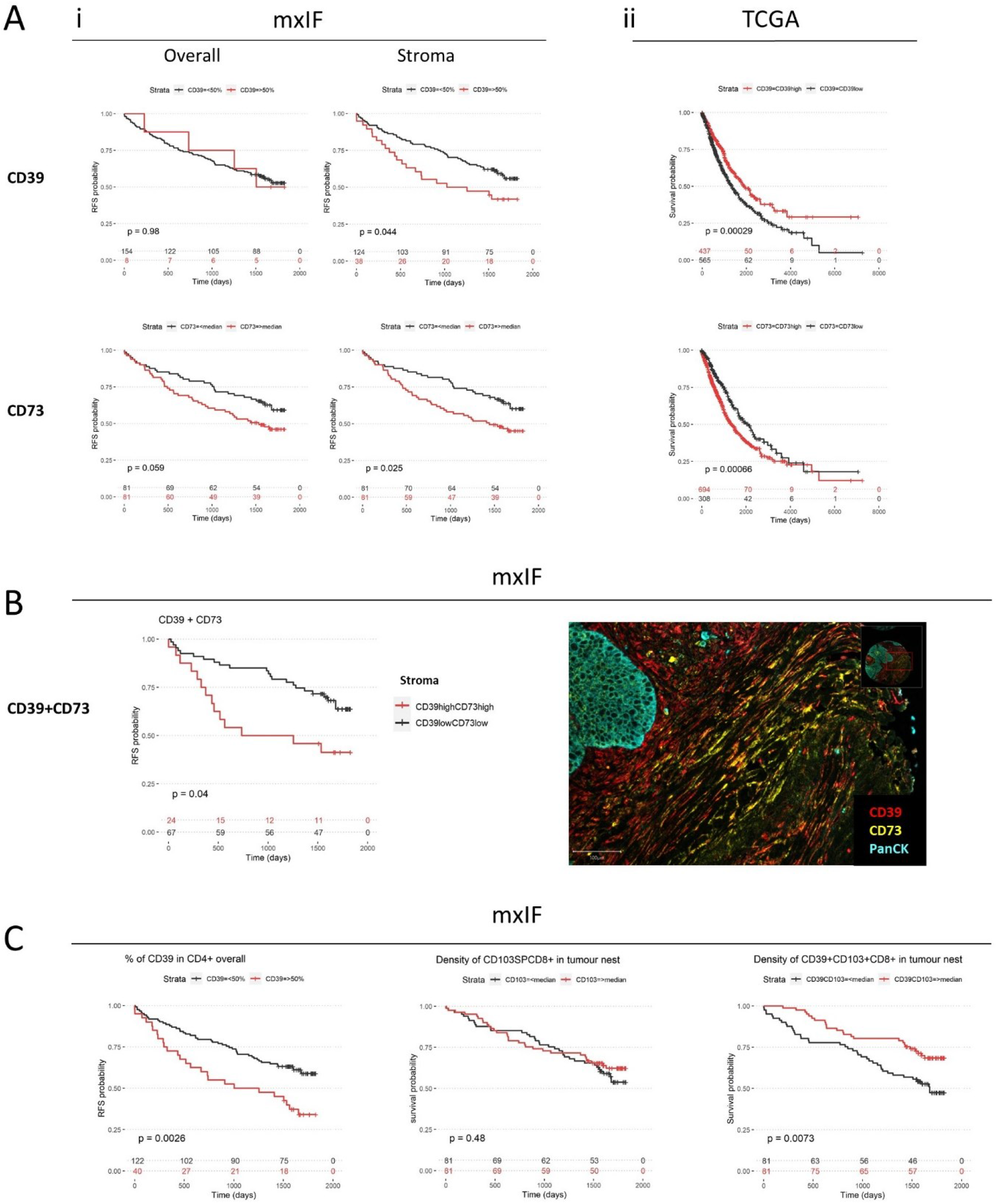
Spatial patterns of distinct T cell populations predict outcome in NSCLC. (A) Kaplan Meier curves showing (i) 5 year RFS based on overall and stromal expression of CD39 (top panel) and CD73 (bottom panel) of mxIF data of early untreated NSCLC pateints (n=162) and (ii) 19 year OS probability based on high/low CD39 (top) and CD73(bottom) of TCGA data. (B) 5 year RFS based on combined stromal CD39 and CD73 expression of mxIF data (left) and representative image of stromal CD39 and CD73 staining in tumour tissue of NSCLC (right). Log Rank tests were used for survival analysis

Using TCGA data, we similarly see that CD73 predicts reduced long-term survival probability, as previously shown by Gao et al.. Interestingly, overall CD39 expression in the TCGA data confers improved long-term survival perhaps due to its high expression levels amongst immune cells and tumour specific CD8+ T cells (Fig.5Aii). To assess this, we investigated predictive value of distinct T cell subpopulations considering their spatial signatures as identified by our immunofluorescence dataset. We see a marked reduction of survival probability in patients with high CD39 expression amongst CD4+ T cells overall and in the stroma, in line with their regulatory features and phenotypic traits of highly suppressive Tregs. (Fig.5C) In stark contrast, the density of CD39+CD103+ CD8+ T cells in the tumour nest significantly predicts favourable survival probability, while the density of bystander CD103SP CD8+ T cells does not (Fig.5C). Notably, the frequency of CD39 single positive CD8+ T cells is too low to conduct robust analysis. This supports our flow cytometry data showing that the majority of CD39+ T cells are CD39+CD103+. These observations were broadly independent of tumour size and smoking status. This illustrates prediction of outcome dependent on CD39 location and cellular subtype. It further demonstrated that functional characteristics of CD39+CD103+ CD8+ T cells (which harbour high cytotoxicity and potential to kill autologous cancer cells in vitro) have a significant biological relevance in the tumour nest in situ which ultimately affects patient outcome in a cohort of early untreated NSCLC patients.

## Discussion

Investigating pathways of immune suppression is essential to develop new targets for immunotherapy. In this study, we report that surface enzymes involved in the adenosine pathway are predictive of poor outcome in early NSCLC. CD39 is expressed by multiple immune- and non-immune cell types in the TME of NSCLC and mainly localised in the stroma. CD39 expression amongst T cells encompasses both CD39+CD103-CD4+ T cells associated with transcriptomic signatures of a highly suppressive regulatory phenotype predictive of poor outcome and exhausted yet highly cytotoxic CD39+CD103+ CD8+ T cells which are able to infiltrate the tumour nest where their presence is linked to improved overall- and recurrence free survival at 5 years in a cohort of early treatment naïve NSCLC patients.

T cell exhaustion is a key feature of CTL within the TME. The exhaustion marker CD39 is highly expressed in terminally exhausted CD8+TIL across solid tumours, including NSCLC, where it allows the distinction of tumour-reactive CTL from bystander CD8+TIL.^3^ Duhen et al. discovered that co-expression of CD39 and CD103 identifies exhausted tissue resident memory cells based on gene transcripts typical of T cell exhaustion a gene signature associated with TRM and reduced expression of T cell recirculation genes shown by transcriptomic analysis of these populations in HNSCC and ovarian cancer.^8^ Here, we report that the TME of NSCLC highly upregulates the CD39 encoding gene ENTPD1 in CD8+ T cells. Flow cytometry analysis shows a shift towards CD39+CD103+ co-expressing CD8+ T cells in tumour tissue compared to NCL, and transcriptomic analysis of single cell RNA Sequencing data demonstrates that these CD39+CD103+ CD8+ T cells are enriched in other exhaustion markers yet display a gene signature indicative of high cytotoxicity and cell killing.^23^ *In vitro*, CD39+CD103+CD8+ T cells are able to kill autologous tumour cells, while negligible tumour cell killing is observed in CD39-CD103+ and CD39-CD103-T cells.^8^ In lung cancer, CD103+CD8+ T cells increase upon immunotherapy and are enriched in responders to anti-PD1.^24^ Similarly, a recent study in breast cancer demonstrates that CD39+CD103+ T cells are tumour specific, predict improved patient outcome in triple negative breast cancer and demonstrate that this cell population is responsive to immune checkpoint inhibition *in vitro*.^25^

We have extended these findings to show that in NSCLC, CD39+ CD8+ T cells are confined to the stroma unless they co-express CD103 in which case they are able to infiltrate the tumour nest. Notably, the density of CD39+CD103+ CD8+ T cells in the tumour nest is linked to improved survival and RFS, while the density of CD103SP CD8+ T cells is not, further implying that CD39+CD103+ CD8+ T cells are the main tumour cell killing population in situ. Interestingly, CD39+CD103+ CD8+ T cells are particularly found in malignancies that are immunogenic and responsive in immunotherapy such as triple-negative breast cancer, melanoma and colon cancer, further suggesting that as they are neo-antigen specific and cytotoxic, they are primed to respond to PD1 blockade and could offer a target to stratify patients for checkpoint inhibition. ^8^

In this study, we also report highly expressed levels of CD39 amongst tumour infiltrating CD4+ T cells via flow cytometry mirroring published reports that have classified this population as particularly suppressive Tregs with a high FOXP3 expression, and potent suppressive effects against CD8+ T cell proliferation and effector function that is stable under inflammatory conditions.^18,26–29^ We extend these findings to show that the majority of CD39+ CD4+ T cells in the TME of NSCLC are confined to the tumour stroma and a high frequency of CD39+CD4+ T cells predicts poor outcome. This is reflective of the divergent pattern of CD103 expression in CD4+ and CD8+ populations. Notably CD103+CD4+TRM are highly reduced in the tumour tissue compared to NCL. The inhibitory capacity of CD39+Tregs is in part caused by the enzymatic activity of CD39 resulting in depletion of available eATP and intracellular accumulation of cAMP in effector T cells as well as downstream adenosine production in the presence of CD73. Functionally, we report an upregulation of LAG3, CTLA4 and TIGIT in line with the exhausted phenotype and gene signatures of regulatory activity of T cell activation, proliferation, and differentiation.

Interestingly, we observed an increased expression of CXCL13 in both CD39+CD4+ T cells and CD39+CD103+ CD8+ T cells. Recently, single-cell analysis has identified CXCL13+ CD8+ T cells as a tumour reactive population correlated to favourable response to ICB.^30^ CXCL13 is a chemoattractant which recruits CXCR5+ B cells. In agreement with this we observe a dramatic increase in B cell frequencies in the TME of NSCLC vs. NCL using flow cytometry, the vast majority of which are CD39+, suggestive of a regulatory B cell phenotype,^31^ and could thereby add to the suppressive environment. However, B cell recruitment has also been associated with TLS formation and a beneficial effect for tumour immunity and responsiveness to checkpoint therapy.^32–35^

Overall, we show that the TME of NSCLC upregulates CD39 expression in multiple cell types including CD56+, CD19+ and CD90+ cells maybe suggesting a shared trigger such as TGFb and hypoxia potentiating CD39 expression.^9^ While CD39 is mainly expressed in stromal areas, CD73 is found in both tumour nest and stroma. This suggests that coordinated activity of CD39 and CD73 is likely to occur in the stroma, resulting in high levels of adenosine in this area, providing a barrier to suppress immune cells prior to their entry into the tumour nest. In support of this, high stromal CD39 and CD73 expression predicts poor RFS at 5 years compared to CD39low CD73low tumours.

There is increasing interest in the role of stromal cells in tumour immunity.^36–39^ Our data illustrates CD39 and CD73 expression on stromal cells with characteristic shape of fibroblasts and which are in close proximity to T cells. We have previously described bidirectional upregulation of CD39 and CD73 elicited by CAF – T cell crosstalk.^21^ CAFs can display an autocrine feedforward loop of adenosine production via CD73 and A2B receptors^40^, and synthesize more adenosine in the presence of activated CD39+ T cells^41^. Thus, this crosstalk could be a potent source of adenosine in the TME.

In conclusion, our data suggests that tumour reactive CD39+CD103+ CD8+ T cells can infiltrate the tumour nest where they confer a survival benefit. However, most CD39+ CD8+ T cells are trapped in the tumour stroma where we see highest frequencies of CD39+CD4+ regulatory T cells and CD73+ stromal cells and therefore likely undergo adenosine mediated immunosuppression. Considering their significant role in the adenosine axis and multiple other immunosuppressive mechanistic roles in the TME, targeting CAFs could simultaneously reduce the adenosine mediated immune suppression and allow redistribution of T cells into the tumour nest (as shown in a preclinical study^42^), perhaps leading to an increased frequency of CD39+CD8+ T cells in the tumour nest which we show here improves survival.

## Funding

This work was supported by Cancer Research UK [CRUK Clinician Scientist Fellowship A24867] to ARA; LK is supported by a GlaxoSmithKline-NPL studentship. LM is supported by EPSRC Centre for Doctoral Training in Medical Imaging [EP/L016559/1].

## Acknowledgements

We are grateful for assistance from CIR Flow Cytometry and Shared University Research Facilities, University of Edinburgh. We would also like to thank all the staff at the department of Thoracic Surgery, Royal infirmary of Edinburgh. We also thank Irene Young and Katie Hamilton for undertaking patient consents.

## Supplementary Data

**Figure S1.**
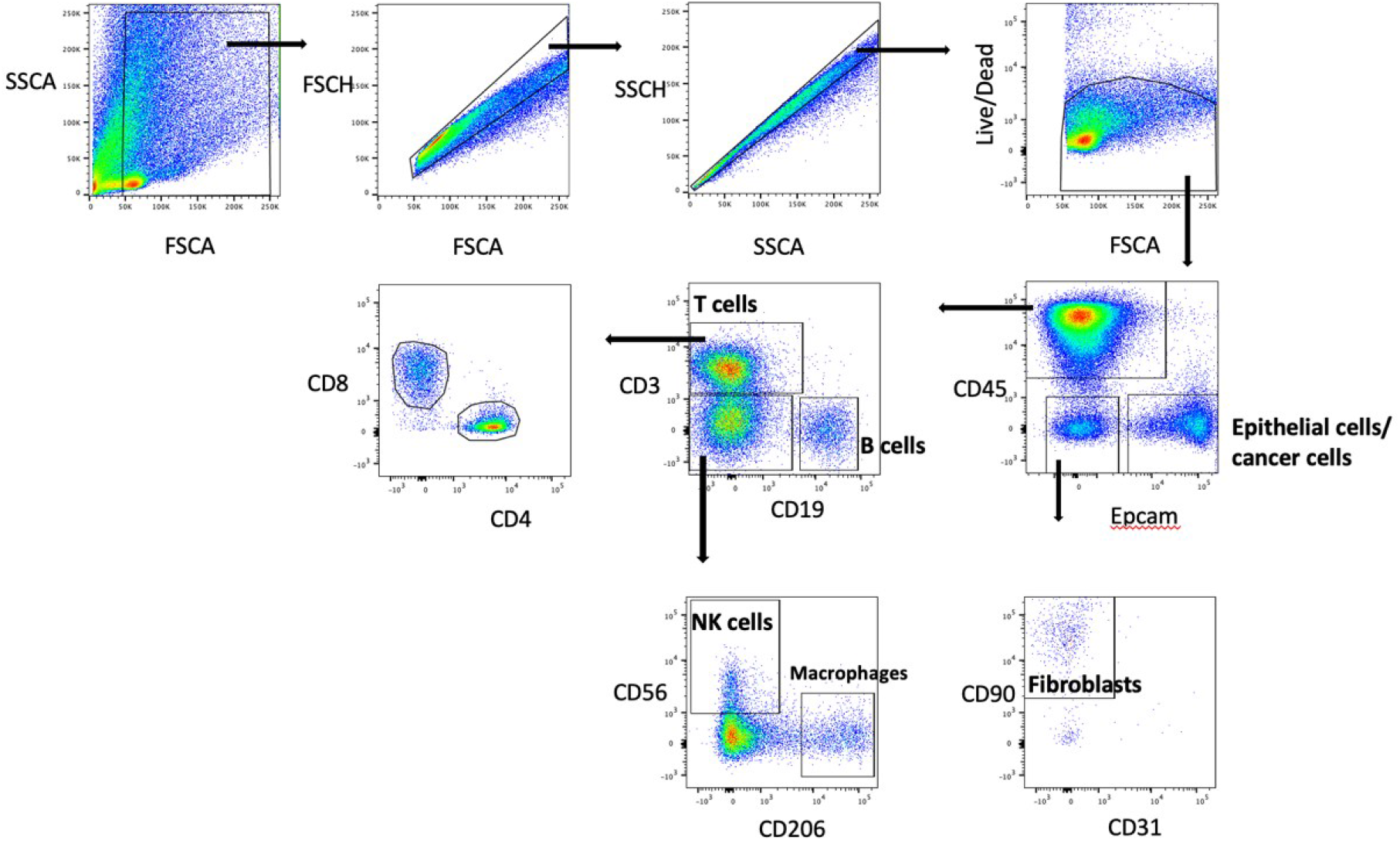
Gating strategy for tissue digest. We first set a cell-gate to remove debris, then gate on single cells, and live cells. Within live single cells, we gate on EPCAM+ and CD45+ cells. Within the Epcam-CD45-population, we gate on CD90+ cells. Within CD45+ cells, we gate on CD19+cells and CD3+cells. Within the CD19-CD3-cells, we gate for NK cells by selecting CD56+ and CD206+. Within CD3+T cells we distinguish between CD4+and CD8+T cells.

**Figure S2.**
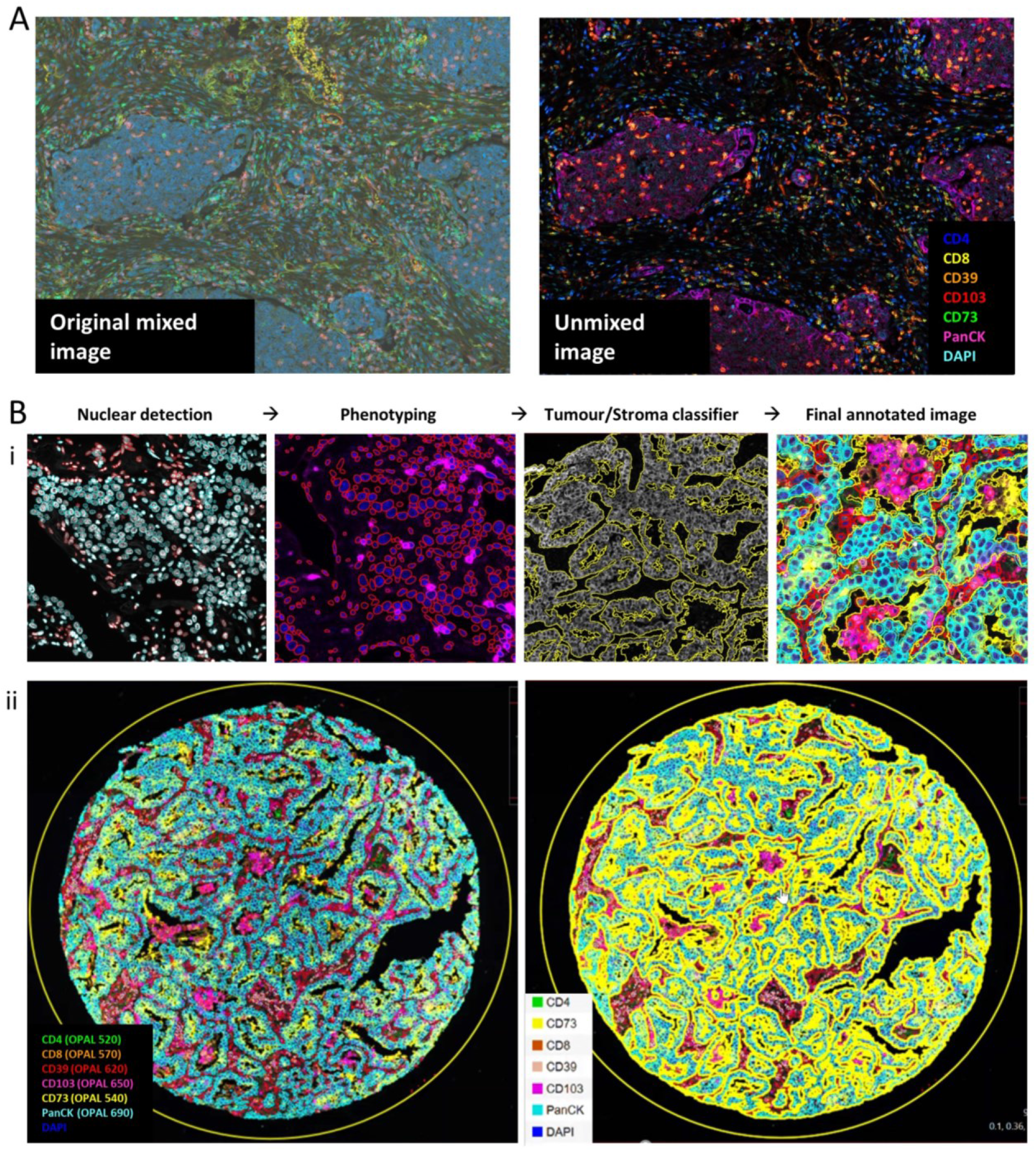
MxIF image analysis workflow. (A) Example image of NSCLC tumour tissue stained with full multiplex panel showing original mixed image (left) and spectrally unmixed image (right). (B) (i) Overview of optimised image analysis workflow in Qupath and (ii) example of original unmixed staining (left) and image showing tumour stroma classified and phenotype annotated image.

**Figure S3.**
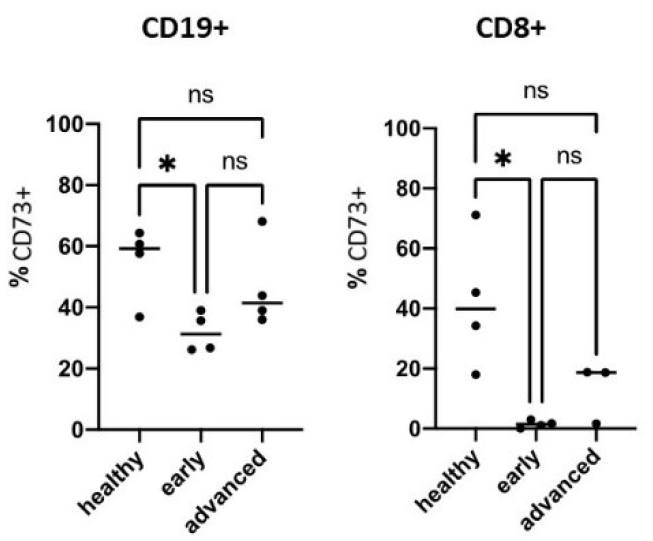
Frequency of CD73+ amongst CD19+ and CD8+ cells isolated from PBMCs of healthy donors and early and advanced NSCLC patients. One way ANOVAs with Tukeys multiple comparisons were used for comparing groups. (*p=<0.05, **p=<0.01, ***p=<0.001)

**Figure S4.**
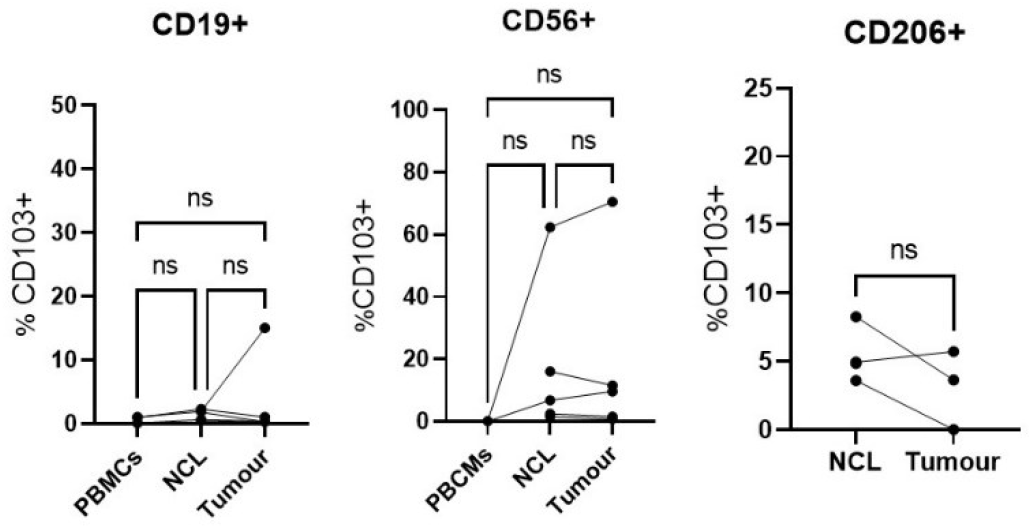
Expression levels of CD103 shown as % positive within subpopulations of CD19+ and CD56+ cells from peripheral blood, NCL and tumour tissue and CD206+ cells from NCL and tumour tissue of early NSCLC patients. Paired t tests were used for statistical analysis of 2 groups, one way ANOVAs with Tukeys multiple comparisons were used for comparing 3 groups. (* p=<0.05, **p=<0.01, ***p=<0.001)

**Figure S5.**
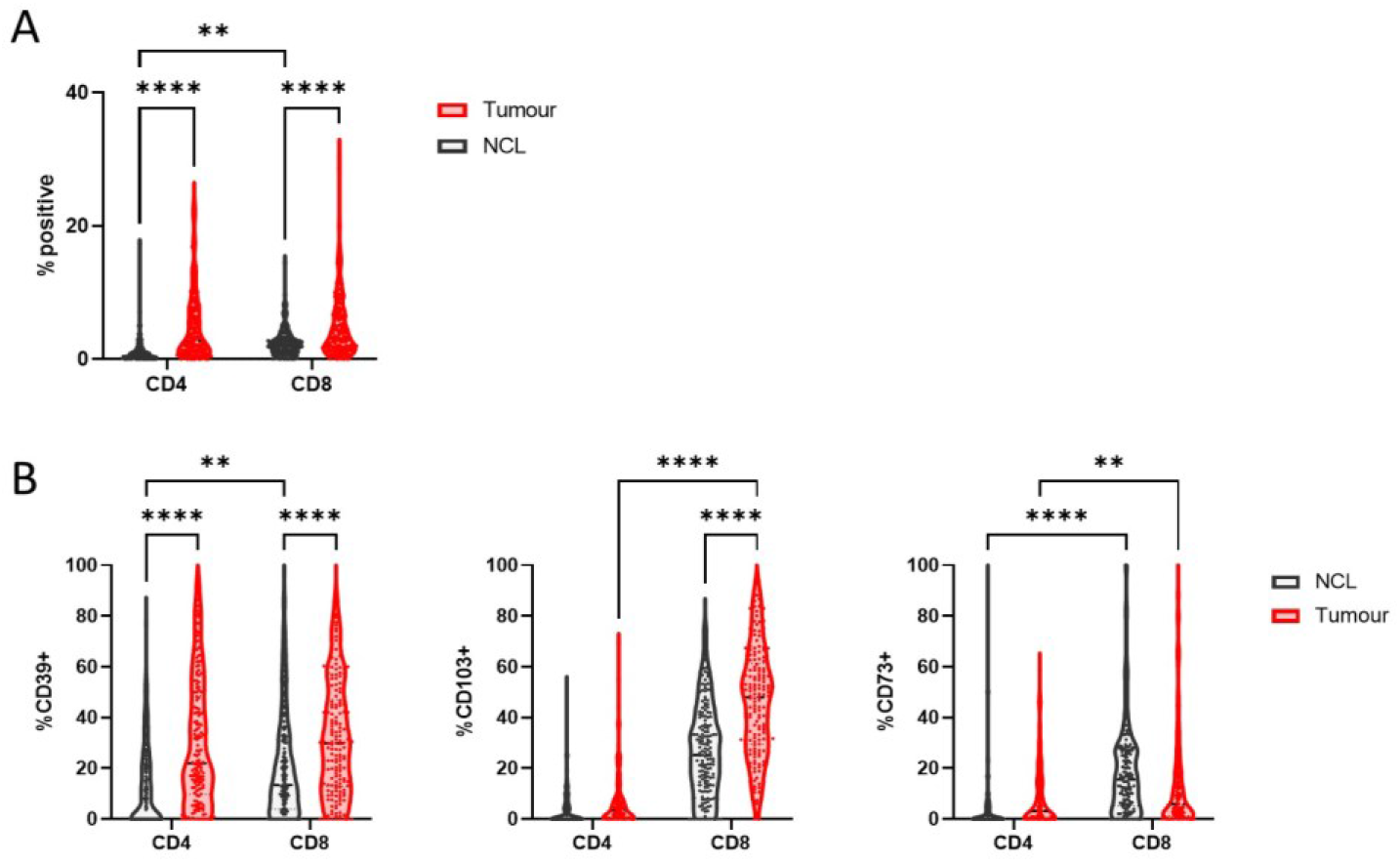
(A) Frequency of CD4+ and CD8+ T cells in tumour compared to NCL tissue. (B) Frequency of CD39+, CD103+ and CD73+ cells within CD4+ and CD8+ T cells in NCL and tumour tissue from MxIF analysis of early untreated NSCLC patients (n=162). Two-way ANOVAs were used to compare groups. (* p=<0.05, ** p=<0.01, ***p=<0.001)

**Figure S6.**
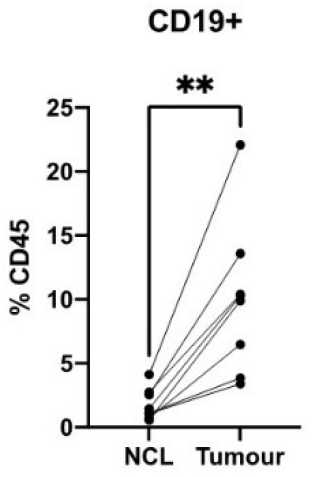
Frequency of CD19+ of CD45+ cells from NCL and tumour tissue of early NSCLC patients. Paired t tests were used for statistical analysis. (* p=<0.05, ** p=<0.01, ***p=<0.001)

**Figure S7.**
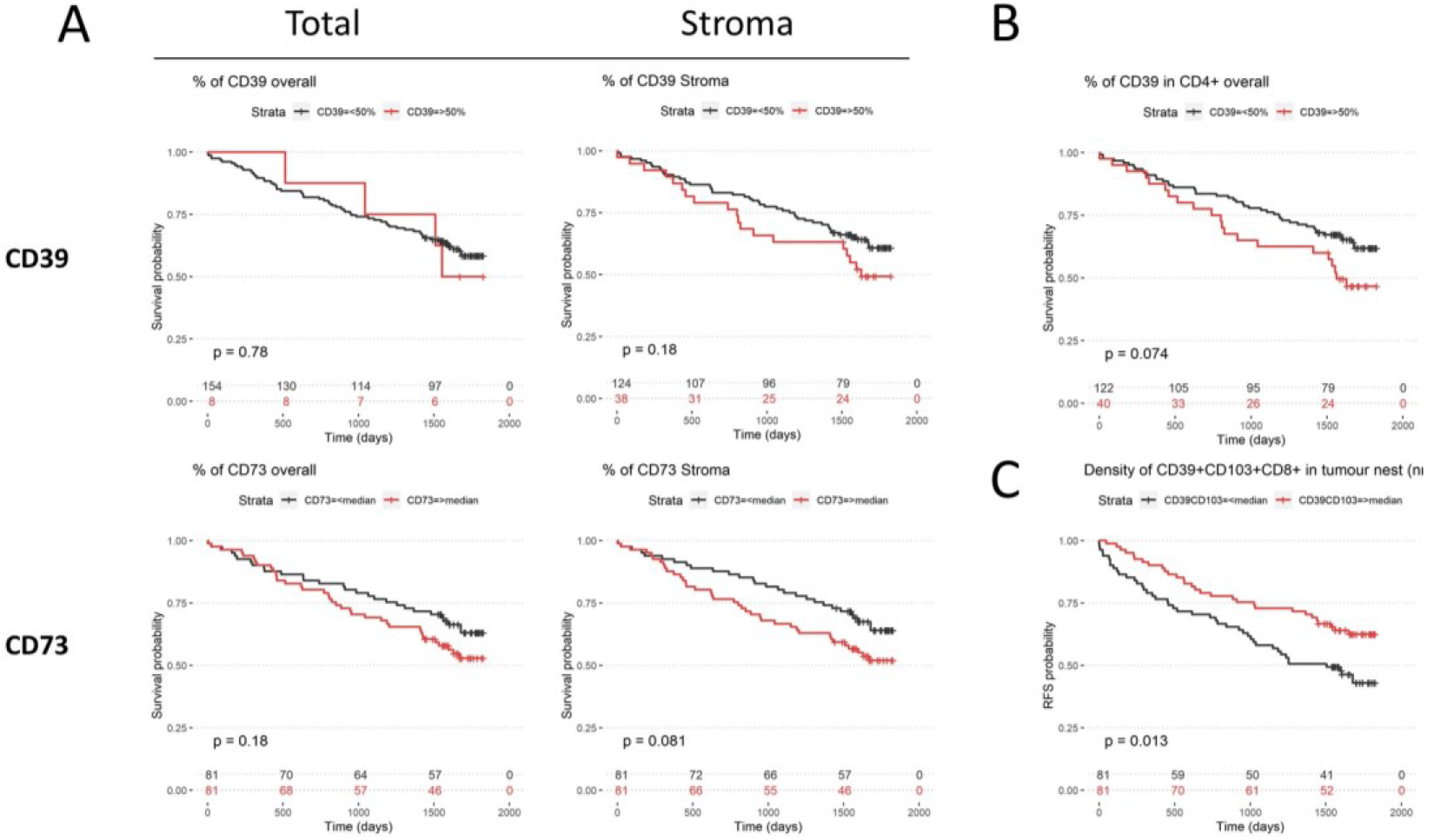
(A) Effect of spatial expression patterns of CD39 (top panel) and CD73 (bottom panel) on survival probability. (B) Effect of frequency of CD39+ amongst CD4+ T cells overall on survival probability and (C) Effect of density of CD39+CD103+ T CD8+ cells on RFS probability at 5 years in early untreated NSCLC patients (n=162). Log Rank tests were used for survival analysis

**Figure S8.**
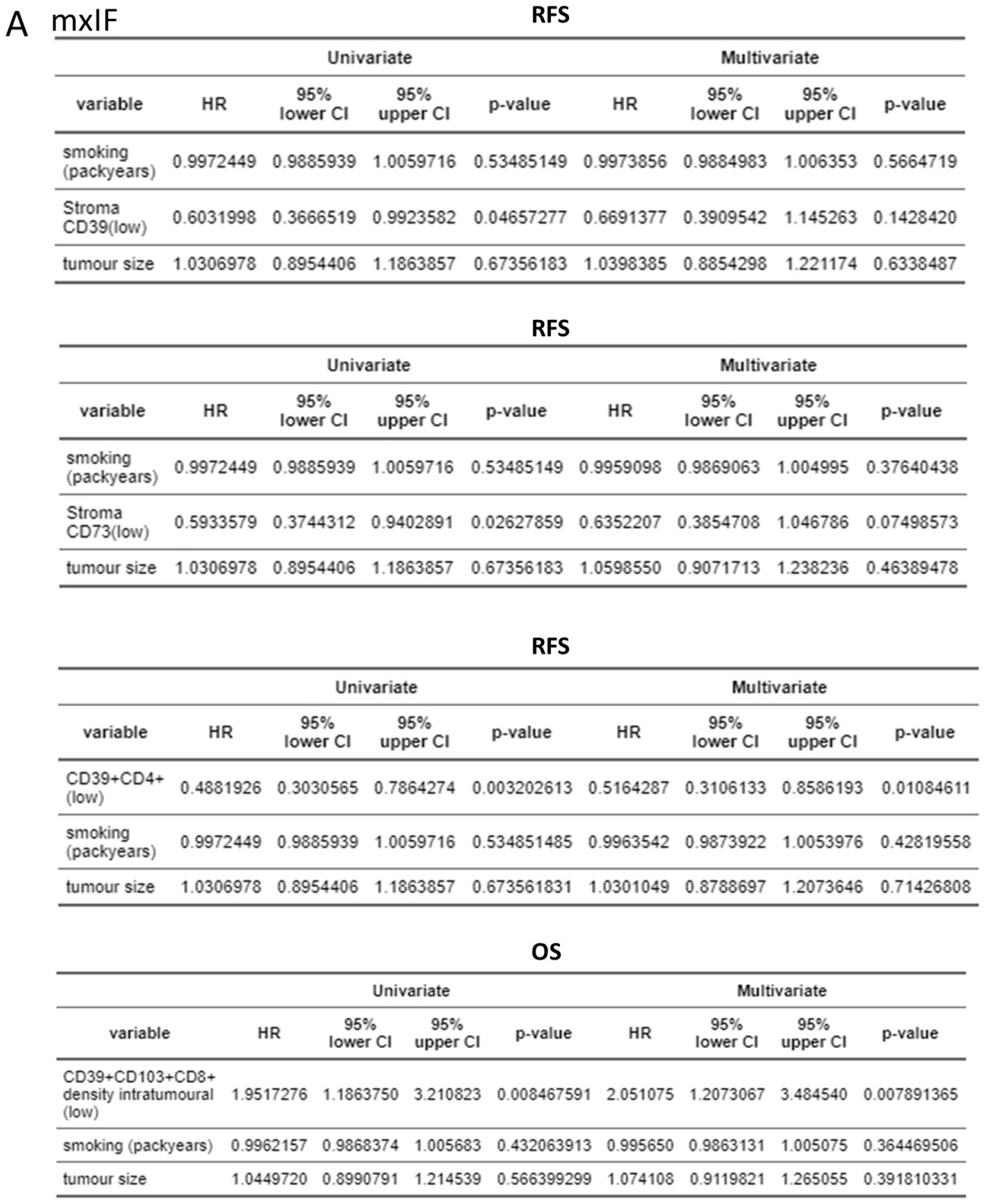
Univariate and multivariate cox regression analysis of smoking, tumour size, and stromal CD39 (</>50% expression)(RFS), stromal CD73 (</> median % expression) (RFS), CD39+CD4+ (</> median % expression)(RFS) and CD39+CD103+CD8+ T cells (density in tumour nest </> median) (OS).

**Figure S9.**
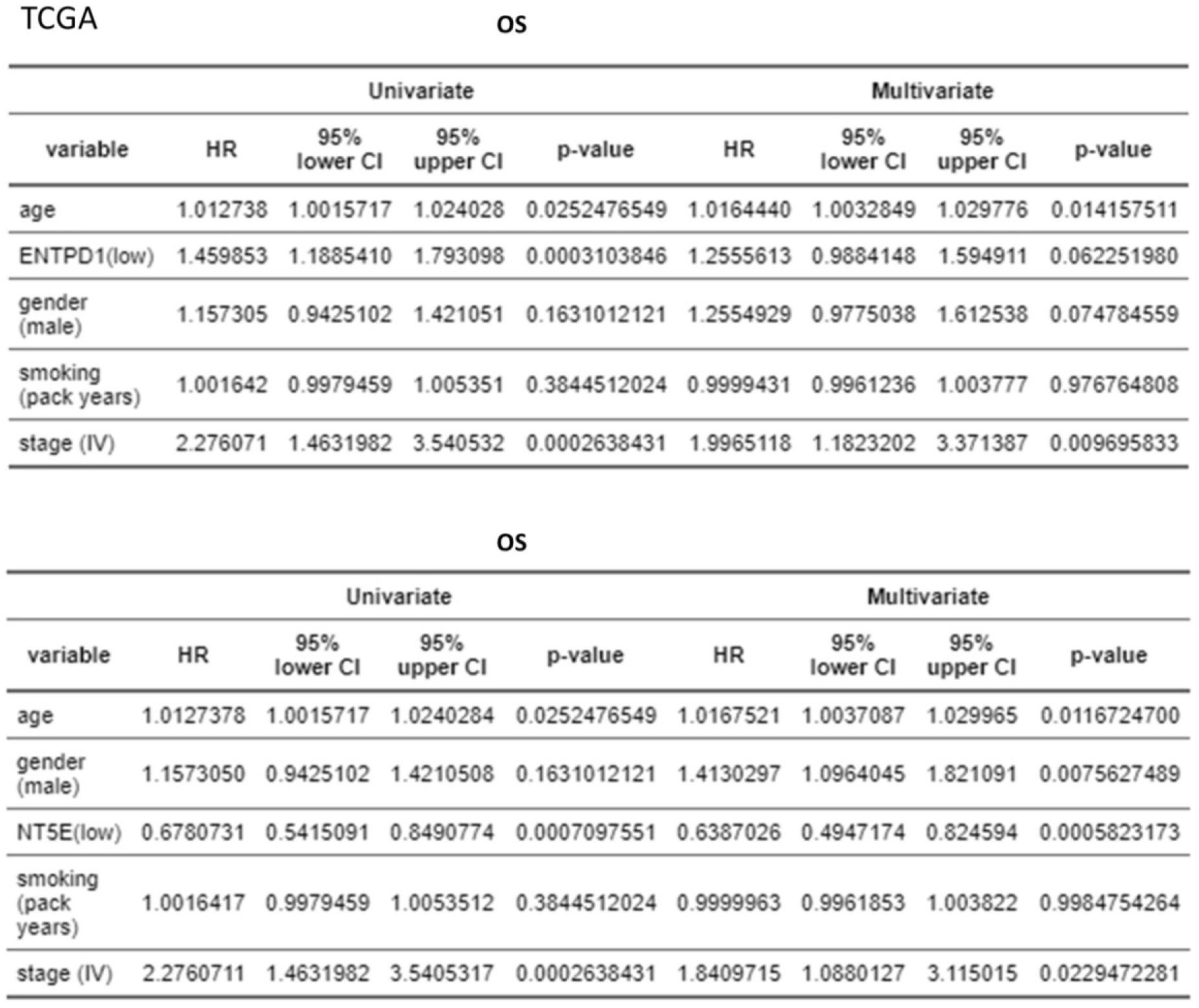
Univariate and multivariate cox regression analysis of OS based on age, gender, smoking, stage and ENTPD1 (low/ high) and NT5E (low/high).

## Notes

### Competing Interest Statement

The authors have declared no competing interest.

